# Nipah virus detection at bat roosts following spillover events in Bangladesh, 2012–2019

**DOI:** 10.1101/2021.12.29.474445

**Authors:** Clifton D. McKee, Ausraful Islam, Mohammed Ziaur Rahman, Salah Uddin Khan, Mahmudur Rahman, Syed M. Satter, Ariful Islam, Claude Kwe Yinda, Jonathan H. Epstein, Peter Daszak, Vincent J. Munster, Peter J. Hudson, Raina K. Plowright, Stephen P. Luby, Emily S. Gurley

## Abstract

Knowledge of the dynamics and genetic diversity of Nipah virus circulating in bats and at the human-animal interface is limited by current sampling efforts, which produce few detections of viral RNA. We report on a series of investigations at bat roosts identified near human Nipah cases in Bangladesh between 2012 and 2019. Pooled bat urine samples were collected from 23 roosts; seven roosts (30%) had at least one sample with Nipah RNA detected from the first visit. In subsequent visits to these seven roosts, RNA was detected in bat urine up to 52 days after the presumed exposure of the human case, although the probability of detection declined rapidly with time. These results suggest that rapidly deployed investigations of Nipah virus shedding from bat roosts near human cases could increase the success of viral sequencing compared to background surveillance and enhance our understanding of Nipah virus ecology and evolution.

## Introduction

Nipah virus is an emerging paramyxovirus (genus *Henipavirus*) that has caused outbreaks of neurologic and respiratory disease in humans and livestock in Bangladesh, India, Malaysia, Singapore, and the Philippines (*1–4*). The primary hosts of henipaviruses are fruit bats (family Pteropodidae) in Africa, Asia, and Australia (*5*). Although Nipah virus causes no apparent disease in bats (*6,7*), the case fatality rate in humans can be 40–70% (*2,8,9*). Additionally, Nipah virus has characteristics that enable repeated outbreaks in humans. Its bat hosts are widespread in South and Southeast Asia, regions with dense human and livestock populations (*10*) that could facilitate virus spillover and spread (*11*). Nipah virus can transmit directly from bats through human consumption of date palm sap contaminated with bat saliva, urine, or feces or indirectly via spillover to domesticated animals (*12–14*). Bangladesh has experienced multiple Nipah outbreaks since 2001 with confirmed person-to-person transmission, albeit below the threshold necessary for sustained epidemics (*8*); however, the virus transmitted rapidly among farmed pig populations in Malaysia, producing infection rates of 100% on some farms, and spread between farms through shipments of infected animals (*15,16*). There are no commercially available vaccines or therapeutics for Nipah virus to prevent or mitigate disease in the case of an epidemic, although this is an area of active research (*17,18*). Finally, Nipah virus is an RNA virus. Although documented genetic diversity within Nipah viruses is limited (*19–23*), the high mutation rates observed in RNA viruses are an important predictor of zoonotic potential (*24*) and could theoretically produce variants with sufficient transmissibility in humans to cause a sustained epidemic (*25,26*). Given the wide geographic range and unsampled diversity of Nipah viruses, it is possible that variants exist and circulate in bats that are more transmissible among humans, and each spillover event is an opportunity for a more transmissible virus to emerge (*27*).

Genetic and phenotypic diversity among Nipah viruses exists, but the human health implications of this variation are unclear. Nipah virus genotypes from Bangladesh and India are genetically distinct from Malaysian genotypes (*21–23*). While the Malaysian genotypes are less diverse than those from Bangladesh and India (*23*), genotypes from Malaysia derive solely from pigs, humans, and bats during the 1998-1999 outbreak whereas genotypes from Bangladesh derive from multiple human outbreaks and surveys of bats since 2004. Another widely cited difference is that person-to-person transmission of Nipah virus was observed rarely in Malaysia (*28–30*) whereas in Bangladesh it accounted for a third of reported cases (*8*) and in India, over 75% of cases (*1,9,31*). However, person-to-person transmission in Malaysia was not widely investigated beyond healthcare workers and less than 10% of Nipah cases transmit the virus to another person, usually a family caregiver (*8,28*). Some of this variation in transmission mode and severity may be attributable to differences in exposure, sampling, infrastructure, and culture between countries, but differences between viral strains may explain additional variation. Cases in Malaysia were less likely to present with a cough or difficult breathing and abnormal chest radiographs than cases in Bangladesh (*29,32,33*). These differences in transmissibility and pathogenicity between Nipah virus strains from Malaysia and Bangladesh have been partly confirmed by animal experiments, although with conflicting results (*34–36*). The reviewed evidence suggests that genetic variation in Nipah virus may produce differences in pathogenicity or transmissibility, so it is possible that more transmissible strains of Nipah virus may circulate undetected in bat populations.

However, our knowledge of Nipah virus diversity is limited to the few virus sequences obtained to date. According to GenBank and recent studies (*19,23*), only 76 Nipah virus genomes and 153 nucleocapsid protein genes have been sequenced, 51 and 37 of which derive from human patients, respectively. Current research efforts are not optimized to characterize Nipah virus genotypes circulating in bats. Longitudinal surveys indicate that exposure to Nipah virus is high (∼40%) in some *Pteropus medius* populations in Bangladesh based on serology, but the prevalence of detectable Nipah virus RNA is low (<5%) at any given time (*37*). In addition, viral loads in collected bat samples are often low (*23*), preventing genetic sequencing or isolation of viruses necessary for describing viral diversity. Sampling methods that increase the success of detecting Nipah virus in bat populations and increase yield so that sequencing is possible would be useful for monitoring genetic changes in this virus. In this study, we focused Nipah virus detection to *Pteropus medius* bat roosts nearby to human cases identified in Bangladesh during outbreak investigations between 2012 and 2019. We aimed to identify whether bat roosts were actively shedding Nipah virus RNA in urine and how long shedding continued after initial detection. Additionally, we sought to identify characteristics of bat roosts that could be associated with higher likelihood of testing positive.

## Materials and Methods

### Nipah virus case investigations

During 2012 to 2019, human cases of suspected Nipah virus infection with a history of date palm sap consumption were identified via surveillance at three hospitals in Faridpur, Rajshahi, and Rangpur Districts in Bangladesh (*38*). Additional suspected cases in other regions were identified year-round from media reports (*39*). Forty-seven primary cases of Nipah virus representing spillover from bats were identified in 2012–2018, of which 17 were investigated in this study. Four additional spillover cases were investigated in 2019, but the total spillovers from that year is unclear due to a lack of reporting. Case exposure to Nipah virus was evaluated with an enzyme-linked immunosorbent assay (ELISA) or PCR (*40*). Investigation teams visited the suspected case villages to gather evidence of case clusters and identify the exposure route (*41*). In some cases, teams were deployed before human cases were confirmed via ELISA or PCR.

Teams searched for *Pteropus medius* bat roosts within a 20 km radius of the case house by asking community members about known roost sites and by scouting. Some identified roosts were located on burial grounds or over water bodies and could not be sampled (Appendix Table 1). Between 12:00 AM and 4:00 AM, teams placed 4 to 20 polyethylene tarps under each roost, depending on the available area and size of the roost, to collect urine. Tarps were concentrated under branches with denser aggregations of bats. Tarps were approximately 6’x4’ in size before 2019 and 3’x2’ in 2019; this change was made so that fewer bats contributed to urine pools to improve estimates of prevalence (*42*). Between 5:00 AM and 6:00 AM, teams returned to the roosts and collected bat urine from the tarps with a sterile syringe. Urine collected from tarps was either pooled by individual tarp or mixed together from multiple tarps and then divided into aliquots. We found no significant difference in Nipah detection between the two strategies (see Appendix). Aliquots were tested for Nipah virus RNA at icddr,b or NIH laboratories using quantitative real-time reverse transcription PCR (RT-qPCR) targeting the nucleoprotein gene (*43*). Roosts with Nipah virus RNA detected in any aliquots at the first sampling event were revisited (3–16 days between sampling events) until all aliquots from a roost tested negative. Attempts to culture Nipah virus from RT-qPCR positive samples at NIH yielded no virus isolates; viral culture was not attempted at icddr,b due to the absence of biosafety level 4 facilities.

### Statistical analysis

For each laboratory-confirmed spillover of Nipah virus in a human, the symptom onset date and the coordinates of the case house were recorded. Teams identified the probable date of patient exposure to Nipah virus via date palm sap consumption for some cases; otherwise, the exposure date was assumed to be seven days prior to symptom onset based on the mean incubation period of Nipah virus for primary cases linked to spillover (*44*).

Logistic regression was used to assess features of the roost sites associated with a roost testing positive for Nipah virus at the first sampling visit. Covariates in the model included the number of days between the first case patient exposure to date palm sap and roost sampling, the number of bats in the roost, the distance between the human case house and the roost site, and the number of human spillover cases associated with each nearby roost. Model selection was then performed to choose important features using Akaike’s corrected information criterion (AICc) (*45*).

For all roost sites that tested positive for Nipah virus at first sampling, we recorded the number of tested urine aliquots that were positive for Nipah virus at each visit. Since cycle threshold (CT) values from RT-qPCR were not reported for all tests, we used the proportion of positive aliquots as an approximate measure of the intensity of virus shedding in roosting bats, assuming that the roosts with higher concentrations of virus in urine will produce more positive aliquots. We then analyzed changes in the proportion of positive aliquots across roosts along two time axes. First, we aligned dates based on the number of days since presumed exposure date of the first human spillover Nipah case associated with each roost site. We also aligned roost sampling dates to the number of days since the start of the calendar year for comparison. Binomial linear models were fit to estimate the probability of detecting a Nipah virus-positive aliquot at each roost along each time axis.

To evaluate the utility of sampling bat roosts nearby to human Nipah cases as a surveillance approach, we compared the rate of successful Nipah virus detections from this study to data recently reported by Epstein et al. (*37*). Samples from that study were collected quarterly from a *P. medius* roost in Faridpur District between 2007 and 2012 as part of a longitudinal study, from visits to different roosts throughout Bangladesh between 2006 and 2011 as part of a cross-sectional spatial analysis, or as part of Nipah outbreak investigations in 2009, 2010, and 2012. Urine samples were either collected from individual bats or from underneath roosts. For these comparisons, we considered each roost visit as a discrete sampling event, including repeat visits to the same roost. Ignoring the initial visits to seven roosts nearby five suspected human cases that were Nipah-negative, the 23 roosts in our study were sampled across 47 visits. Comparisons between studies were made for the number of sampling visits with positive Nipah detections and the number of positive urine samples (individual or pooled aliquots from roosts) across all sampling visits or during the first visit to each roost. Comparisons were evaluated using a chi-squared test of proportions or Fisher’s exact test, where appropriate. Statistical tests were considered significant if p-values were <0.05.

### Ethical approval

All study participants or proxies provided informed consent before participation and personally identifiable information from case patients was delinked from the data before use. Written permission was obtained from the Bangladesh Forest Department for sampling the bats and team members obtained permission from landowners before sampling roosts. Protocols for case investigations and roost sampling were reviewed and approved by the institutional review board at icddr,b.

## Results

Teams investigated roosts near 21 suspected human cases of Nipah virus infection during 2012–2019 (Appendix Table 1). The cases were clustered in the central and northwest districts of Bangladesh, close to the three surveillance hospitals (Figure 1). Symptom onset for patients occurred in winter (December–February) with the exception of one case patient in Manikganj District who presented with symptoms in March 2013. No roost investigations were performed in 2017 and 2018 due funding constraints.

**Figure 1.**
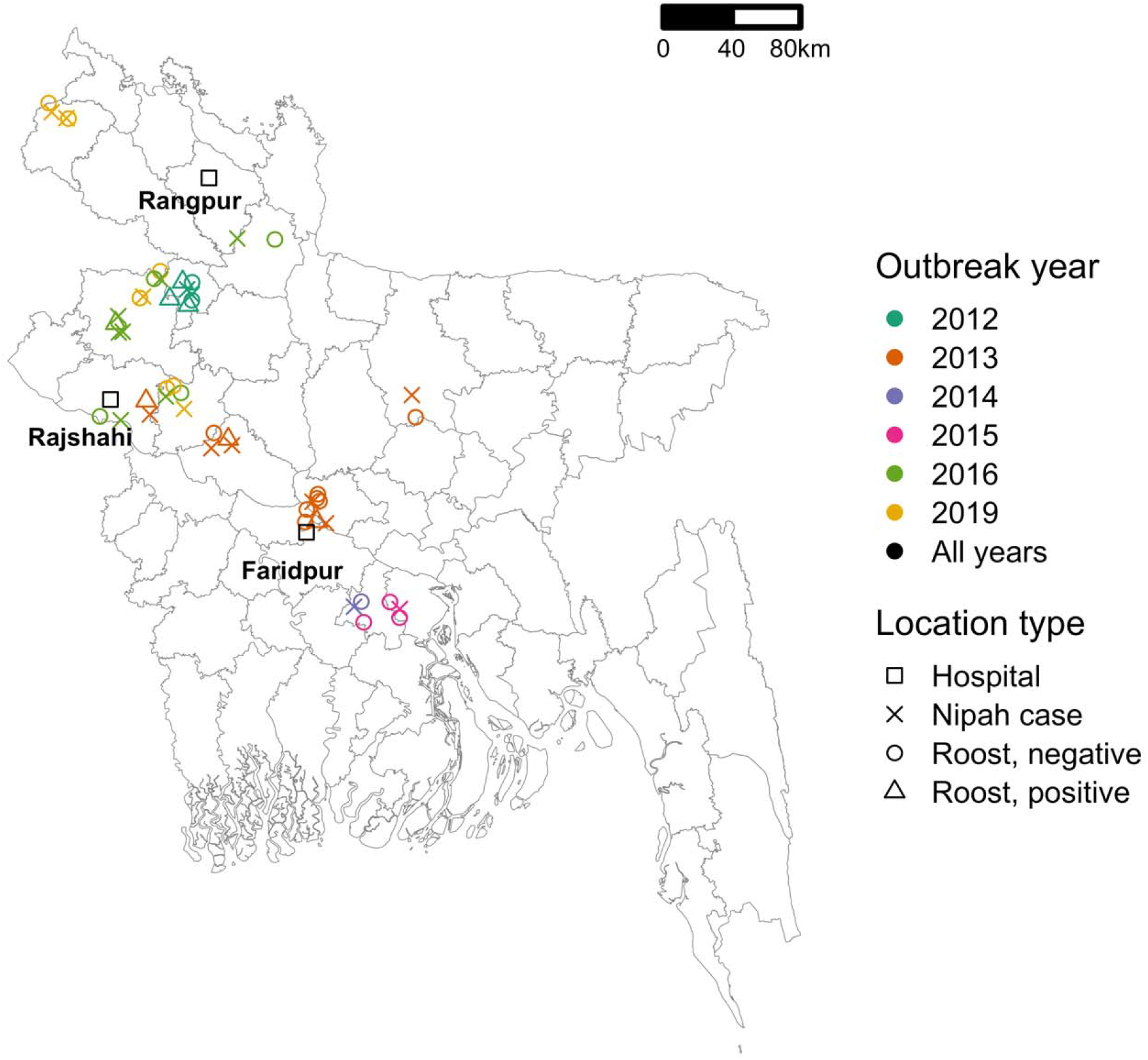
Locations of human Nipah cases (n = 21) and *Pteropus medius* bat roosts (n = 30) investigated in Bangladesh between 2012 and 2019. Roosts with urine aliquots that tested positive for Nipah virus RNA at the first sampling visit are marked with distinct symbols. Points have been jittered a small amount to increase visibility.

For each case patient, 1–3 *Pteropus medius* roosts were identified within 0–17.9 km of the patient’s home (Appendix Dataset 1). Five additional roosts were identified but could not be sampled due to their location on burial grounds or over water (Appendix Table 1). A total of 30 roosts were sampled. The first sampling visits occurred 17–62 days after the case patients’ exposure to date palm sap, either reported from the case investigation or back-calculated as seven days prior to the onset of symptoms (Appendix Dataset 1). Five of the suspected patients tested negative for Nipah virus by ELISA or PCR, and the seven roosts identified near the case houses yielded no Nipah virus RNA. Since our interest was in whether sampling around Nipah cases would help to identify roosts with active Nipah virus shedding, the suspected but Nipah test-negative case patients and associated bat roosts were left out of statistical analyses. Sensitivity analyses with these samples included produced statistically similar results. RT-qPCR testing of pooled urine aliquots detected 7/23 (30%) roosts as positive for Nipah virus RNA in one or more aliquots at the first sampling visit.

Logistic regression on the presence of Nipah virus RNA in roost urine at the first sampling event was performed on 22 distinct roosts using four explanatory variables; one roost was omitted due to missing data on the number of bats. Although roosts with positive urine aliquots tended to have more associated human Nipah spillover cases, were sampled sooner after patient exposure, were more distant from the case house, and had a smaller number of bats, none of these variables was significantly associated with roost positivity in univariate or multiple regression analyses (Figure 2; Appendix Table 2) and AICc identified the intercept-only model as the best model (Appendix Table 3).

**Figure 2.**
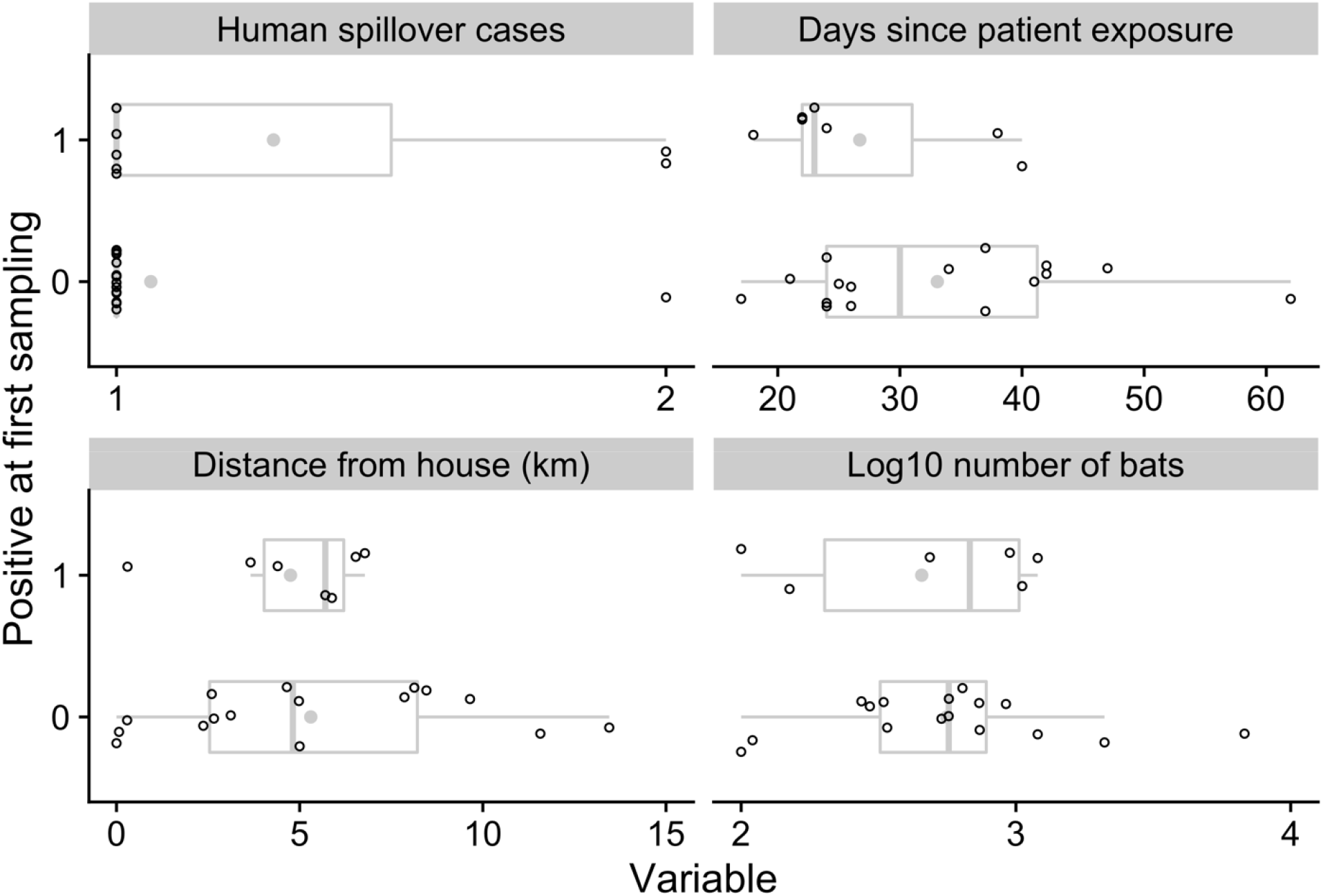
Descriptive variables for 23 *Pteropus medius* roosts sampled nearby confirmed human Nipah virus cases. Open points show the values associated with the first human case associated with each roost. Means for each variable and positivity status (0 or 1) are displayed as filled gray points.

For the seven roosts where NiV RNA was detected at least once, data were compiled on the number of urine aliquots that tested positive at each repeated sampling visit. Four of the seven roosts were positive at the first visit only and were subsequently only revisited once. The other three roosts remained positive at 1–2 additional sampling visits, although the proportion of aliquots that tested positive declined rapidly with the time since exposure of the first associated human case (Figure 3). For the two roosts with reported CT values from RT-qPCR, the proportion of positive aliquots decreased over the repeated sampling visits while CT values increased, suggesting a decline in viral load with time (Appendix Table 4).

**Figure 3.**
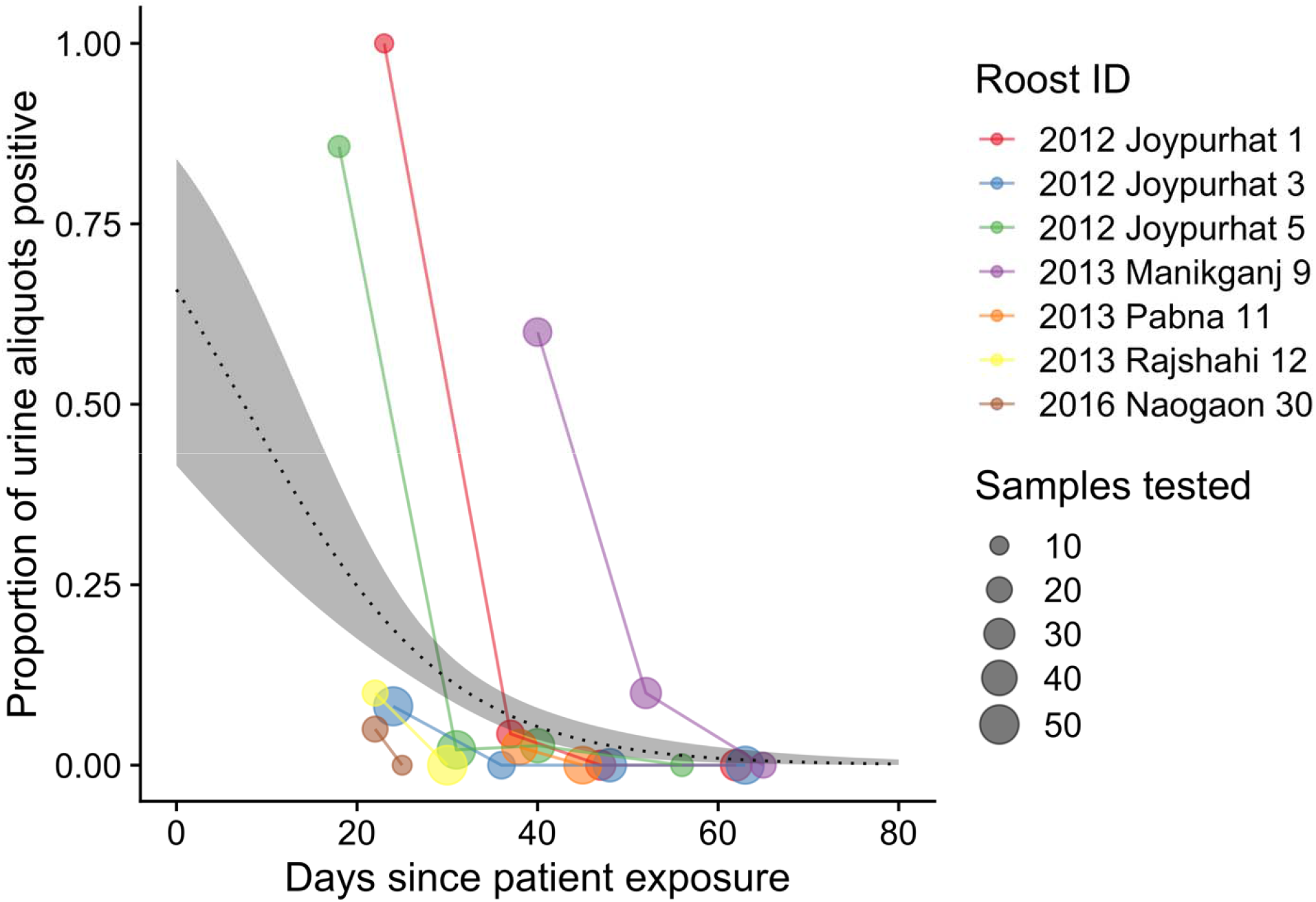
Results of screening *Pteropus medius* roost urine aliquots for Nipah virus RNA. For each roost, the proportion of urine aliquots out of the total tested (shown by the size of points) is aligned along a time axis of the days since the first associated case patient was exposed to Nipah virus in date palm sap, which was either reported during the investigation or back-calculated as five days prior to reported symptom onset.

Fitting a binomial model to the PCR data predicted that the probability of detecting at least one urine aliquot from under-roost sampling as positive for Nipah virus RNA at the time (day 0) the associated case patient was presumably exposed was 0.66 (95% confidence interval (CI): 0.42–0.84) (Figure 3). This probability declined to 0.02 (95% CI: 0.01–0.04) by day 52 when the last positive roost aliquots were detected and to 0.01 (95% CI: 0–0.02) by day 65 when the last roost was sampled. We also fit a binomial model using the days elapsed since the start of the calendar year (Appendix Figure 1), but the alignment of the virus detections among the roosts was less clustered on this time axis than the days since patient exposure time axis, and the binomial model did not show a significant trend in virus detection over time.

Roost urine samples from the current study and individual urine samples from longitudinally sampled roosts in the Epstein et al. study (*37*) produced similar proportions of positive sampling visits (Table 1, comparison A); the detection rate was also similar if only the first visit to each roost in our study was considered (7/23, 30%). In contrast, the proportion of positive aliquots from all sampling visits was significantly higher in our investigations than in the individual urine samples from longitudinal roosts in Epstein et al. (Table 1, comparison B). The detection rate from our study for positive urine aliquots at the first sampling visit was also higher than the detection rate for individual urine samples collected from eight roosts from a cross-sectional study by Epstein et al. (Table 1, comparison C). The detection rate for positive urine aliquots from our study was substantially higher than the detection rate from similarly pooled urine aliquots from underneath longitudinal and cross-sectional roosts in the Epstein et al. study (Table 1, comparison D). Lastly, outbreak investigations of roosts performed by Epstein et al. produced a higher detection rate than our own roost investigations (Table 1, comparison E), although only four roosts were visited by Epstein et al. and the same roosts were not repeatedly visited as was done in our study.

**Table 1.**
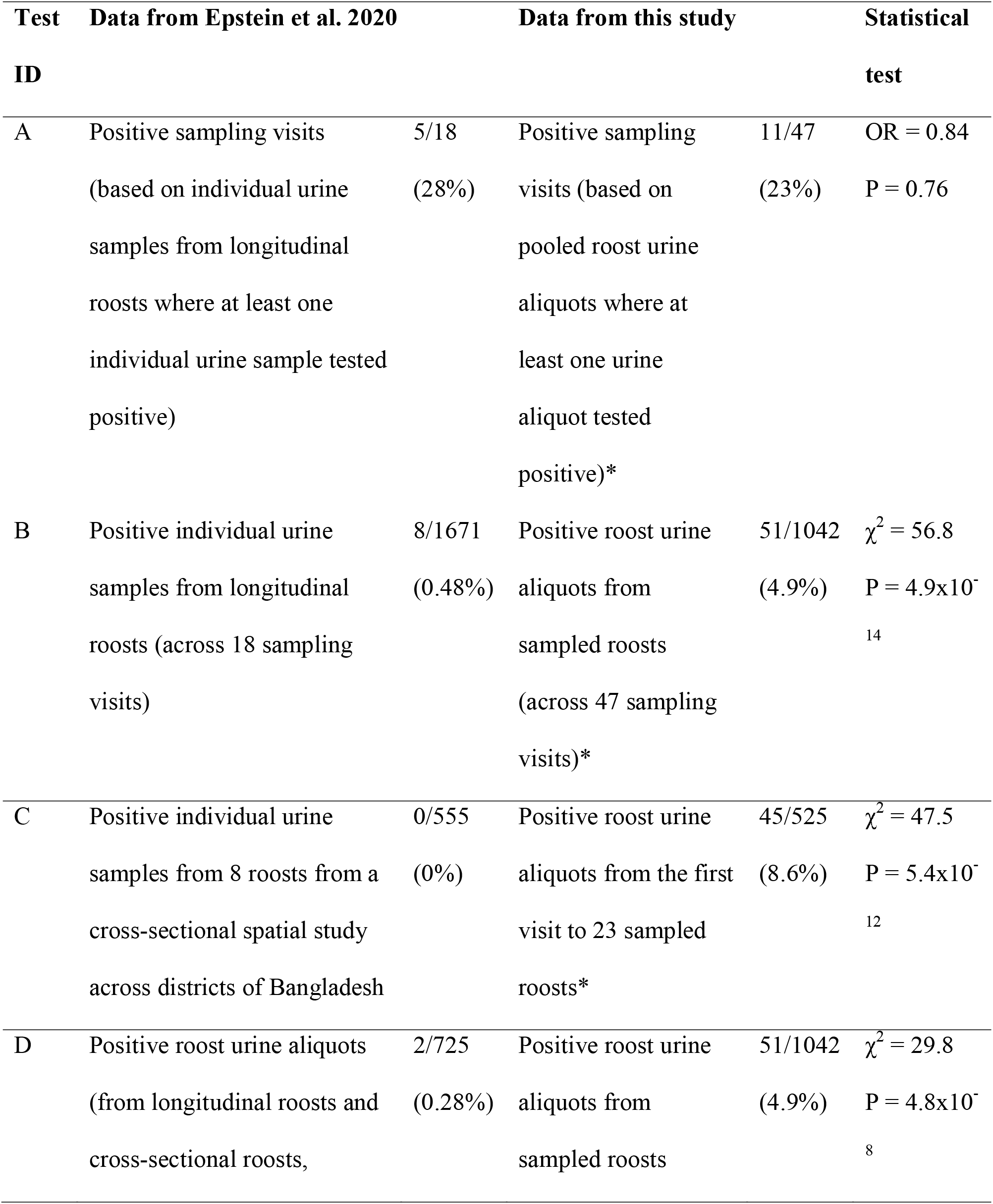
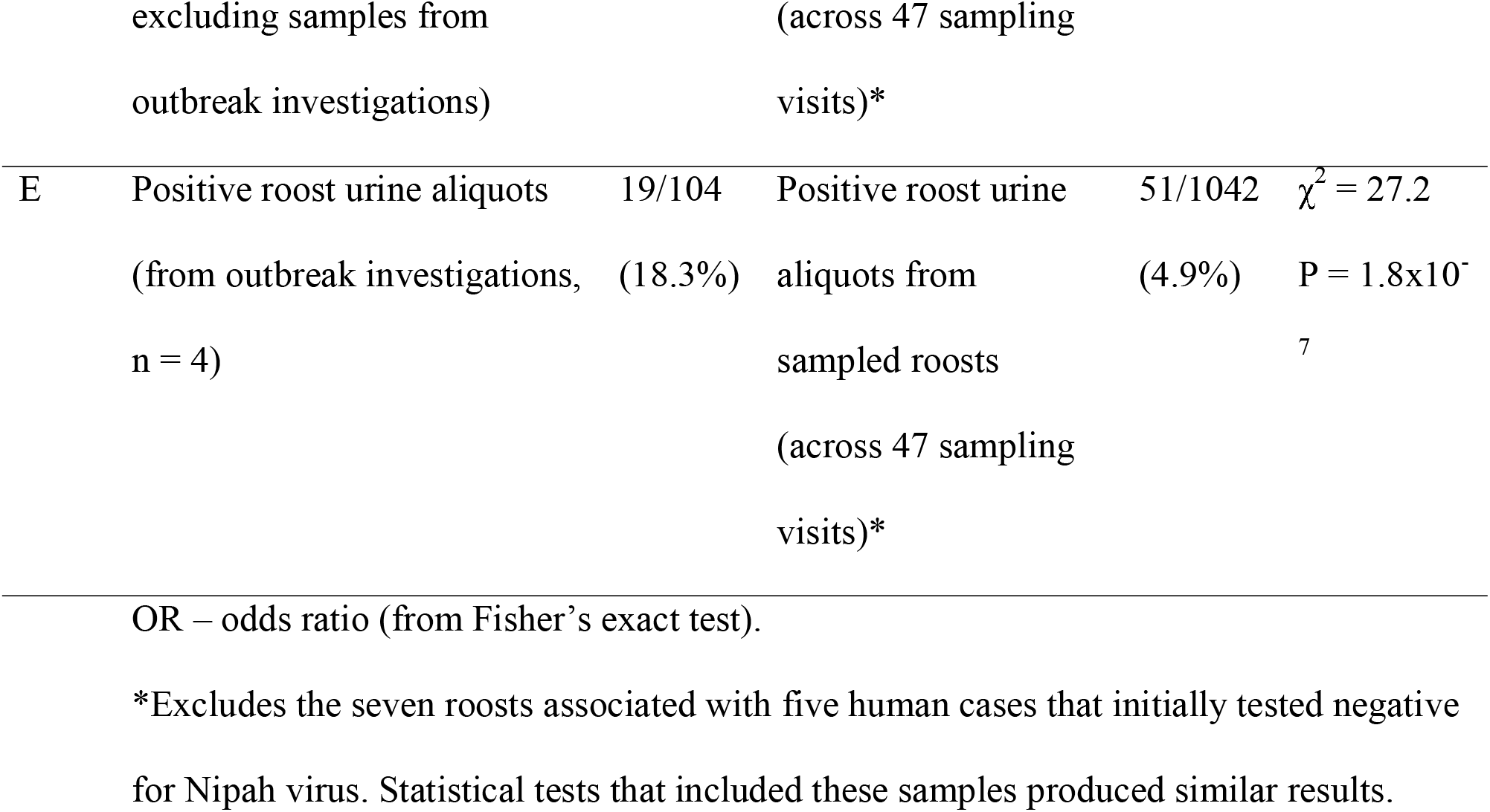
Comparison of Nipah virus detection success from this study with results from Epstein et al. 2020 (*37*).

## Discussion

Nipah virus spillovers from bats occur sporadically in Bangladesh, so surveillance that optimizes viral detection from bat populations is a challenge. In contrast with cross-sectional or longitudinal roost surveillance used previously in Bangladesh (*37*), the roost sampling in this study was triggered by Nipah outbreaks in nearby villages. Our approach identified bat roosts with active Nipah virus shedding at an equivalent rate to background surveillance (*37*), but had a higher detection rate in roost urine on a per sample basis. These results indicate that investigating roosts near spillover cases is more efficient than cross-sectional or longitudinal surveillance for obtaining samples with detectable viral RNA (Table 1). Repeated visits to positive roosts also demonstrated that viral RNA was detectable for weeks after the purported exposure date of human cases, although the proportion of positive urine aliquots declined sharply over time. These data suggest that rapid deployment of teams to identify bat roosts and sample urine could increase the probability of detecting and sequencing Nipah virus. Used in combination with longitudinal sampling of bat roosts and surveillance of human or domesticated animal cases, this method could enhance our understanding of Nipah virus dynamics and genetic diversity in bat populations.

Furthermore, this study provides important knowledge about the timing of Nipah virus shedding in bats in Bangladesh. Longitudinal surveys have shown that Nipah virus shedding from bats is sporadic throughout the year (*37*), so the peaks in viral detection in roost urine from our study likely coincided with shedding events. However, because these shedding events were occurring during winter when date palm sap is harvested for human consumption, bat visits to date palm trees to consume sap may be more likely to contaminate sap with virus and lead to human infections (*46*). This suggests that the intensity of shedding events in bats occurring in winter could help to explain some of the spatial and temporal variation in the number of human spillovers that occur in Bangladesh each year (*41*), although more data on the frequency and timing of shedding events and human sap consumption patterns will be needed to fully understand the dynamics of Nipah virus spillover.

Our findings come with several caveats due to limitations in our sample size and the study design. Our analysis of factors associated with a roost testing positive at the first sampling visit was unable to identify any significant relationships likely due to low statistical power. We also did not systematically attempt Nipah virus isolation or sequencing in all positive samples, so we cannot estimate the probability of successful isolation or sequencing. However, Nipah virus isolates and sequences have been obtained from some of the roost urine samples included this study. One of the positive roosts sampled in Joypurhat in 2012 produced nine nucleocapsid sequences (GenBank accession numbers: MT890702-MT890710; Rahman et al. (*23*)) and the positive roost in Manikganj from 2013 produced ten virus isolates with full-genome sequences (GenBank accession numbers: MK575060-MK575069; Anderson et al. (*20*)). In fact, of the 39 Nipah virus sequences obtained from bats in Bangladesh, 28 (72%) came from urine collected underneath roosts and 24 (86%) of those urine samples came from roost investigations near human cases (Appendix Dataset 2). These patterns suggest that roost urine, especially from roosts near human spillover cases, may contain sufficient Nipah virus for sequencing or culture. Future investigations could track how viral load in roost urine varies during viral shedding events, which could improve the success of sequencing and isolation and shed light on the ecological and epidemiological conditions that lead to Nipah shedding from bats (*47*).

Our case investigations were also limited to the catchment area of three surveillance hospitals and to the winter seasonality of Nipah spillover surveillance. This design systematically misses virus shedding events at bat roosts outside of the surveillance area or during seasons when people are not drinking fresh date palm sap (*13*). Acknowledging that the logistical constrains of our surveillance approach cannot capture all Nipah virus genotypes circulating in *P. medius* across Bangladesh, increasing the number of detections is still important, especially given the few Nipah isolates currently available (n = 11). Reactive roost investigations could be complemented with additional roost surveys outside of surveillance areas to understand Nipah virus transmission and genetic diversity in bat populations across Bangladesh.

This study provides proof of concept that reactive investigations of bat roosts nearby human Nipah cases can complement ongoing surveillance efforts and could increase the likelihood of viral detection and sequencing. Improvements in virus detection would aid in characterizing the genetic diversity of Nipah viruses circulating in bats and identify novel genotypes that might pose a pandemic threat. Furthermore, these data provide evidence that viral shedding can continue for weeks after an initial spillover event, posing a hazard for additional contamination. Knowledge of the period over which bats are shedding Nipah virus could be used to deploy public health campaigns more efficiently, such as using barriers to prevent bat access to date palm sap (*48*).

## Acknowledgments

We acknowledge the Bangladesh Forest Department, the Ministry of Environment and Forest for their permission to conduct these investigations. We thank Robert Fischer and Trenton Bushmaker for technical assistance with bat sample screening. This work was funded by the DARPA PREEMPT program Cooperative Agreement (D18AC00031). Additional funds came from the National Institutes of Health (NIH) grant number 00991 and National Academy of Science (NAS) grant number PGA-2000002048. CKY and VJM are supported by the Intramural Research Program of the National Institute of Allergy and Infectious Diseases, National Institutes of Health (1ZIAAI001179-01). RKP was supported by the US National Science Foundation (DEB-1716698) and the USDA National Institute of Food and Agriculture (Hatch project 1015891). icddr,b acknowledges with gratitude the commitment of NIH, NAS, and DARPA to its research efforts. icddr,b is also grateful to the Governments of Bangladesh, Canada, Sweden, and the UK for providing core/unrestricted support. The views, opinions and/or findings expressed are those of the authors and should not be interpreted as representing the official views or policies of the Department of Defense or the US Government.

## Author Bio

Dr. McKee is a postdoctoral fellow in the Department of Epidemiology, John Hopkins Bloomberg School of Public Health. His primary research interests include microbiology, epidemiology, and wildlife disease ecology.

Dr. Islam is an assistant scientist in the Infectious Diseases Division at icddr,b. His primary research interests include zoonotic disease ecology and epidemiology.

---

**Appendix Table 1.** *Pteropus medius* roosts identified and sampled near suspected human index cases of Nipah virus infection in Bangladesh, 2012–2019. Roosts that were identified but not sampled were not accessible because they were located on burial grounds or over water.

| Year | Division | District | Suspected Nipah cases | Total identified roosts | Sampled roosts | Unsampled roosts |
| --- | --- | --- | --- | --- | --- | --- |
| 2012 | Rajshahi | Joypurhat | 2 | 6 | 5 | 1 |
| 2013 | Mymensingh | Mymensingh | 1 | 1 | 1 | 0 |
|  | Dhaka | Manikganj | 1 | 3 | 3 | 0 |
|  | Rajshahi | Pabna | 2 | 2 | 2 | 0 |
|  |  | Rajshahi | 1 | 1 | 1 | 0 |
| 2014 | Dhaka | Madaripur | 1 | 1 | 1 | 0 |
|  |  | Manikganj | 1 | 3 | 3 | 0 |
| 2015 | Dhaka | Madaripur | 1 | 3 | 3 | 0 |
| 2016 | Rajshahi | Joypurhat | 1 | 1 | 1 | 0 |
|  |  | Naogaon | 3 | 1 | 1 | 0 |
|  |  | Natore | 1 | 1 | 1 | 0 |
|  |  | Rajshahi | 1 | 1 | 1 | 0 |
|  | Rangpur | Gaibandha | 1 | 1 | 1 | 0 |
| 2019 | Rajshahi | Naogaon | 1 | 2 | 2 | 0 |
|  |  | Natore | 1 | 5 | 2 | 3 |
|  |  | Rajshahi | 1 | 2 | 1 | 1 |
|  | Rangpur | Thakurgaon | 1 | 1 | 1 | 0 |

**Appendix Table 2.**
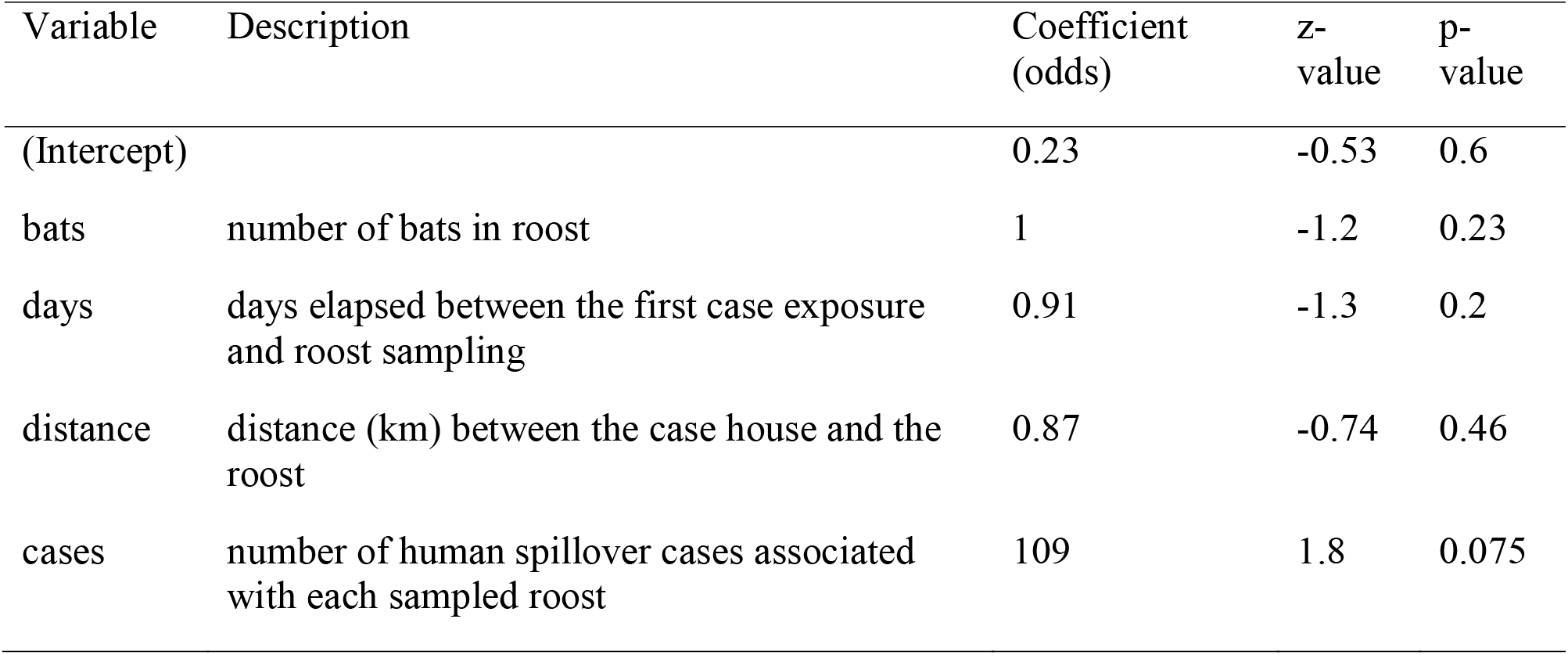
Multivariate logistic regression model coefficients for the presence of Nipah virus RNA at *Pteropus medius* roosts (n = 22) near human cases.

| Variable | Description | Coefficient<br>(odds) | z-<br>value | p-<br>value |
| --- | --- | --- | --- | --- |
| (Intercept) |  | 0.23 | -0.53 | 0.6 |
| bats | number of bats in roost | 1 | -1.2 | 0.23 |
| days | days elapsed between the first case exposure<br>and roost sampling | 0.91 | -1.3 | 0.2 |
| distance | distance (km) between the case house and the<br>roost | 0.87 | -0.74 | 0.46 |
| cases | number of human spillover cases associated<br>with each sampled roost | 109 | 1.8 | 0.075 |

**Appendix Table 3.** Model selection for variables associated with the presence of Nipah virus RNA at *Pteropus medius* roosts (n = 22) near human cases.

| Model | df | AICc | ΔAICc |
| --- | --- | --- | --- |
| (Intercept) | 1 | 27.98 | 0 |
| cases | 2 | 28.01 | 0.03 |
| bats + cases | 3 | 28.33 | 0.35 |
| cases + days | 3 | 28.78 | 0.8 |
| days | 2 | 29.02 | 1.04 |
| bats + cases + days | 4 | 29.5 | 1.51 |
| bats | 2 | 30.03 | 2.05 |
| distance | 2 | 30.4 | 2.42 |
| cases + distance | 3 | 30.67 | 2.69 |
| bats + cases + distance | 4 | 31.17 | 3.19 |
| bats + days | 3 | 31.26 | 3.28 |
| cases + days + distance | 4 | 31.5 | 3.57 |
| days + distance | 3 | 31.71 | 3.73 |
| bats + cases + days + distance | 5 | 32.3 | 4.32 |
| bats + distance | 3 | 32.73 | 4.75 |
| bats + days + distance | 4 | 34.16 | 6.18 |

**Appendix Table 4.** Changes in the proportion of urine aliquots testing positive over repeated visits and associated cycle threshold (CT) values from RT-qPCR.

| Roost | Sampling date | Urine aliquots positive | Average CT value |
| --- | --- | --- | --- |
| 2013 Manikganj 9 | 11 April 2013 | 15/25 (60%) | 30.5 |
|  | 23 April 2013 | 3/30 (10%) | 36.8 |
|  | 6 May 2013 | 0/20 (0%) | ND |
| 2013 Rajashahi 12 | 1 February 2013 | 2/20 (10%) | 38.6 |
|  | 9 February 2013 | 0/50 (0%) | ND |

**Appendix Figure 1.**
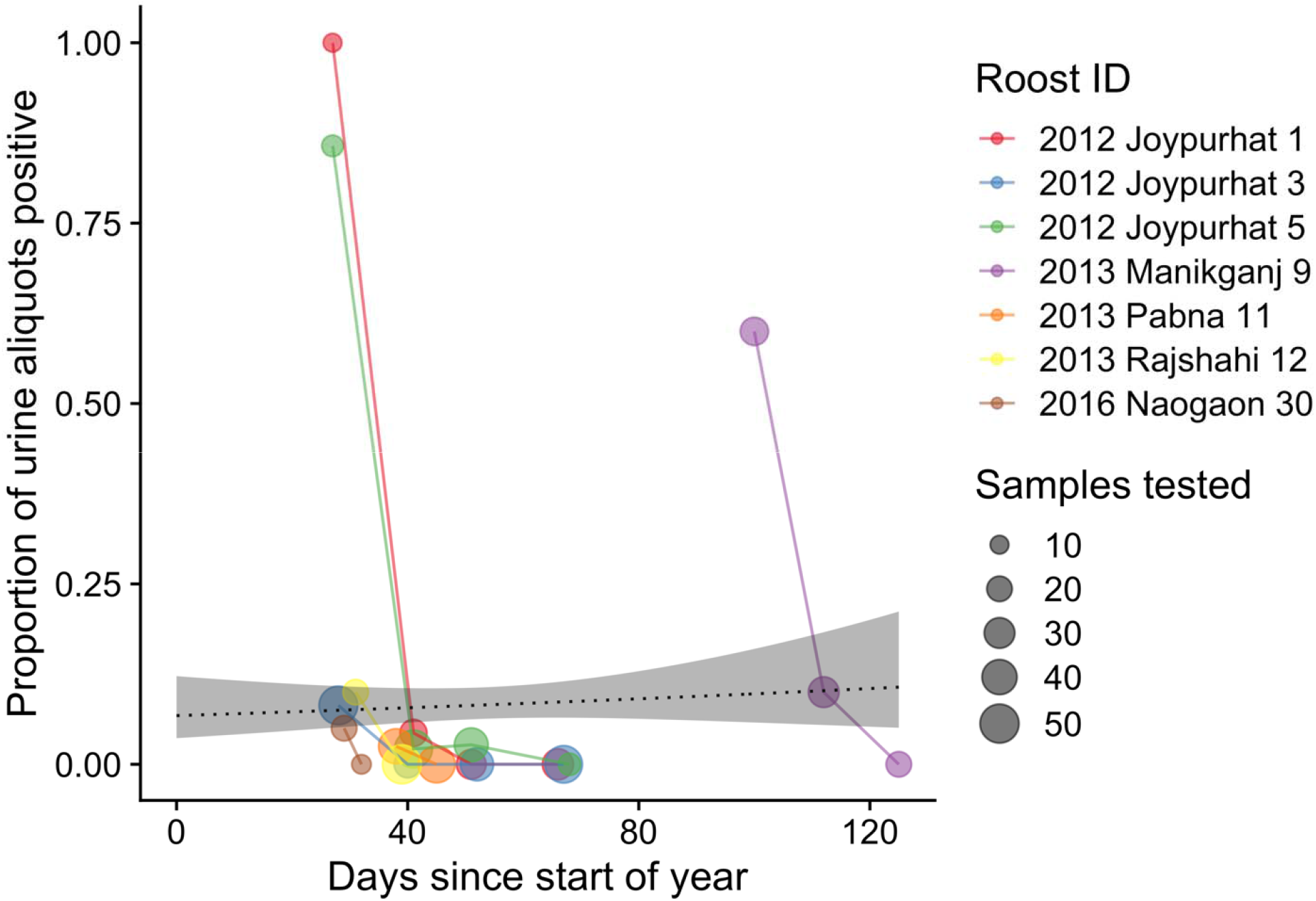
Results of screening *Pteropus medius* roost urine aliquots for Nipah virus RNA. For each roost, the proportion of urine aliquots out of the total tested (shown by the size of points) is aligned along a time axis of the days since the start of the calendar year for each roost investigation.

**Appendix Dataset 1**. Data on the location of roosts, testing dates, and status of human cases.

**Appendix Dataset 2**. Compilation of publicly available Nipah virus sequences obtained from bats and human cases.

