## Appendix Text for "Nipah virus detection at bat roosts following spillover events in Bangladesh, 2012–2019"

**Supplementary Materials and Methods**

*Pooling of roost urine*

Collected urine from underneath roosts was aggregated in 50 ml Falcon tubes from tarps either individually (i.e., one tube per tarp) or mixed together from all tarps and then divided into aliquots for testing: 0.25–0.3 ml of urine in 0.75–1.0 ml of viral transport medium (VTM) or lysis buffer prior to 2014, or equal volumes (0.1–0.3 ml) of urine and media after 2014. The two tarp pooling strategies were used at different sampling events throughout the period of the study. According to our records, 22 sampling events pooled urine from all tarps together and 24 events pooled urine individually by tarp (the pooling strategy for one event could not be determined from field records). For three sampling events where urine was pooled individually by tarp, multiple aliquots from different tarps were positive for Nipah RNA by PCR. Additionally, for the sampling events where urine was pooled from all tarps, the proportion of PCR positive aliquots ranged from 0–100%. We did not observe a clear pattern that indicated that the large pooled volume of urine in these sampling events strictly prevented viral detection due to dilution of viral RNA (i.e., all aliquots were negative or only a few were positive) or that aggregating urine across pools led to saturation (i.e., all aliquots were positive). We could not detect a significant difference between the two pooling strategies in terms of the proportion of positive sampling events: 7/22 (32%) for events with all tarps mixed together vs. 4/24 (17%) for events with separate pools per tarp (Fisher’s test odds ratio = 1.89, P = 0.5). We also did not detect a significant difference between the strategies in terms of the proportion of positive aliquots across all sampling events: 30/527 (5.7%) for events with all tarps mixed together vs. 21/499 (4.2%) for events with separate pools per tarp (χ^2^ = 0.9, P = 0.34). Therefore, we concluded that these differences in pooling strategy did not appear to interfere with our ability to detect Nipah RNA in roost urine and were suitable to be analyzed together.
